# Neuronal activation in the axolotl brain promotes spinal cord regeneration

**DOI:** 10.1101/2024.12.19.629459

**Authors:** SE Walker, K Yu, S Burgess, K Echeverri

**Affiliations:** Marine Biological Laboratory, Eugene Bell Center for Regenerative Biology, Woods Hole, MA; Translational and Functional Genomics Branch, National Human Genome Research Institute, National Institutes of Health, Bethesda, MD

## Abstract

The axolotl retains a remarkable capacity for regenerative repair and is one of the few vertebrate species capable of regenerating its brain and spinal cord after injury. To date, studies investigating axolotl spinal cord regeneration have placed particular emphasis on understanding how cells immediately adjacent to the injury site respond to damage to promote regenerative repair. How neurons outside of this immediate injury site respond to an injury remains unknown. Here, we identify a population of dpErk^+^/etv1^+^ glutamatergic neurons in the axolotl telencephalon that are activated in response to tail amputation. Importantly, the activation of these neurons facilitates successful tail regeneration by promoting axon extension into the newly regenerated tissue. We demonstrate that these dpErk^+^ neurons extend axons into the hypothalamus, which has a well-established role in hormone production. Furthermore, these neurons upregulate the neuropeptide neurotensin in response to injury, which in turn stimulates the production of multiple hormones in the hypothalamus to promote regenerative repair. Together, these findings identify a unique population of neurons in the axolotl brain whose activation is necessary for successful spinal cord regeneration, and sheds light on how neurons outside of the immediate injury site respond to a spinal cord injury.

## Introduction

Regeneration is a widespread phenomenon that is found across the animal kingdom, ranging from whole-body regeneration in flatworms ^1–9^ to liver regeneration in humans ^10,11^. Although many animals are capable of regenerative repair in some capacity, very few species can regenerate their central nervous system (CNS), including the brain and spinal cord, after a traumatic injury ^12–17^.

The axolotl, *Ambystoma mexicanum*, has a remarkable capacity for regeneration, and is one of the few vertebrate species capable of regenerating its brain and spinal cord ^18–22^. In mammalian systems, a traumatic injury to the spinal cord results in Wallerian degeneration, in which damaged neurons surrounding the injury site degenerate ^23–29^. In addition to this widespread neuronal death, glial cells rapidly migrate to the lesion site to form a physical barrier around the injury, known as the glial scar. While the glial scar is thought to prevent further damage to the spinal cord, it also acts as a physical barrier that obstructs regenerating axons from extending into and across the lesion site, ultimately preventing functional regeneration ^23–30^. Unlike these mammalian systems, the axolotl does not form glial scar tissue, and its neurons possess the intrinsic capacity to regenerate after a traumatic injury, ultimately reestablishing lost synaptic connections to promote functional recovery ^22,31^.

In the past two decades, work in the axolotl has largely focused on how cells immediately adjacent to the injury site promote functional spinal cord regeneration, and has uncovered multiple signaling pathways that are necessary to prevent the formation of glial scar tissue ^19,31–34^. However, the spinal cord is innervated by long-range projection neurons that often lie vast distances away from the injury site, such as the brain and brainstem ^22,35^. Importantly, such long-range projection neurons are critical for regulating voluntary movement and are therefore necessary for functional spinal cord repair ^36^. Thus, if we are to fully elucidate how neurons in regeneration-competent species are able to successfully regenerate in response to a traumatic spinal cord injury, we must also investigate neuronal responses that lie outside of the immediate injury site.

In primates, long-range projecting corticospinal tract neurons that regulate voluntary movement of the lower extremities are born in the motor cortex of the brain, where they form monosynaptic contacts with motor neurons in the lower spinal cord ^37,38^. In addition to these corticospinal tracts, primates also house rubrospinal and reticulospinal circuits, in which neurons born in the cortex of the brain synapse with interneurons in the midbrain and brainstem, which in turn extend into the spinal cord. Together, these descending circuits work in concert to regulate voluntary movement ^35,39^. Indeed, the deletion of corticospinal tract neurons in primates leads to paralysis in the lower extremities, however, this can be moderately restored over time ^40^. A more extensive deletion of corticospinal tract neurons in addition to either reticulospinal or rubrospinal circuits, however, leads to permanent paralysis ^41,42^. While corticospinal, rubrospinal and reticulospinal descending circuits are important for voluntary function in primates, adult rodents appear to lack corticospinal tracts. Instead, they house rubrospinal and reticulospinal circuits that regulate voluntary movement ^43^. However, regardless of the exact architecture of these circuits, long-range projection neurons that reside in the brain and/or brainstem are important for voluntary movement. Similar to these vertebrate species, the axolotl contains long-range projection neurons that extend from the brain into the spinal cord ^22^, yet overall, the axolotl brain circuits remain largely unmapped. However, no studies to date have examined how these long-range circuits respond to an injury in the lower spinal cord.

Classical work carried out in other salamander species has revealed a role for signaling from the pituitary gland, including the production of growth hormone to be essential for limb regeneration ^44–47^. In axolotls, limb amputation leads to the activation of cell division in tissues in other parts of the animal; this effect has been attributed to adrenergic signaling ^48^. Collectively, these data suggest that there is a long-range signaling system between distal injuries and the brain.

Here, we identify a population of neurons in the medial pallium of the axolotl telencephalon that are activated in response to spinal cord injury. Inhibiting the activity of these cells impairs normal spinal cord regeneration, corresponding to shorter tail regenerate lengths and fewer axons extending into the newly regenerate tail tissue. We further demonstrate that these neurons express the mammalian marker, *etv1*, which is expressed in neurons that project from the cortex of the brain to the spinal cord. Interestingly, despite the expression of *etv1*, viral tracing demonstrated that these cells primarily extend into the hypothalamus of the brain rather than the spinal cord. Single-cell sequencing analyses determined that these neurons upregulate the neuropeptide *neurotensin* (*nts)* in response to injury. We discovered inhibiting *nts* after injury leads to defects in tail regeneration, and impairs the production of hormones in the brain that may promote regenerative repair. Collectively, this work demonstrates that a distinct population of glutamatergic *etv1*^+^ neurons in the axolotl brain are activated in response to injury, and that their activation is necessary to promote spinal cord repair.

## Results

### Injury leads to long-lasting Erk activation in the brain

Extracellular-regulated kinase (Erk) signaling has become a well-characterized signaling pathway that is required for a variety of neuronal processes, including proliferation, differentiation, and synaptogenesis ^49–56^. After its dual phosphorylation, Erk is activated and in turn phosphorylates multiple downstream targets, including transcription factors that activate gene expression. Thus, a nuclear dpErk localization has become a well-established marker for neuronal activation. We have previously identified Erk as a gene that is upregulated in glial cells within the spinal cord after a spinal cord ablation injury ^57^, but its role in long-range neuronal signaling in the axolotl remains unknown.

To determine whether neurons outside of the immediate injury site are activated in response to a spinal cord injury, we stained the axolotl brain with a diphosphorylated Erk (dpErk) antibody at various time-points after tail amputation. In uninjured axolotls, minimal dpErk signal was detected in the brain (**Fig. 1A**). However, as early as 30-minutes post tail amputation, we discovered a robust increase in Erk signaling in the axolotl brain (**Fig. 1B**). This increase in dpErk expression was located in the cell bodies of neurons and appeared to be a long-lasting response that was detected until approximately 43-days post tail amputation (**Fig. 1B-F**). At 43-days post tail amputation, some Erk signal was still detected in the brain; however, this did not correspond to neuronal cell bodies but was instead detected at low levels in the cytoplasm (**Fig. 1F**).

**Figure 1.**
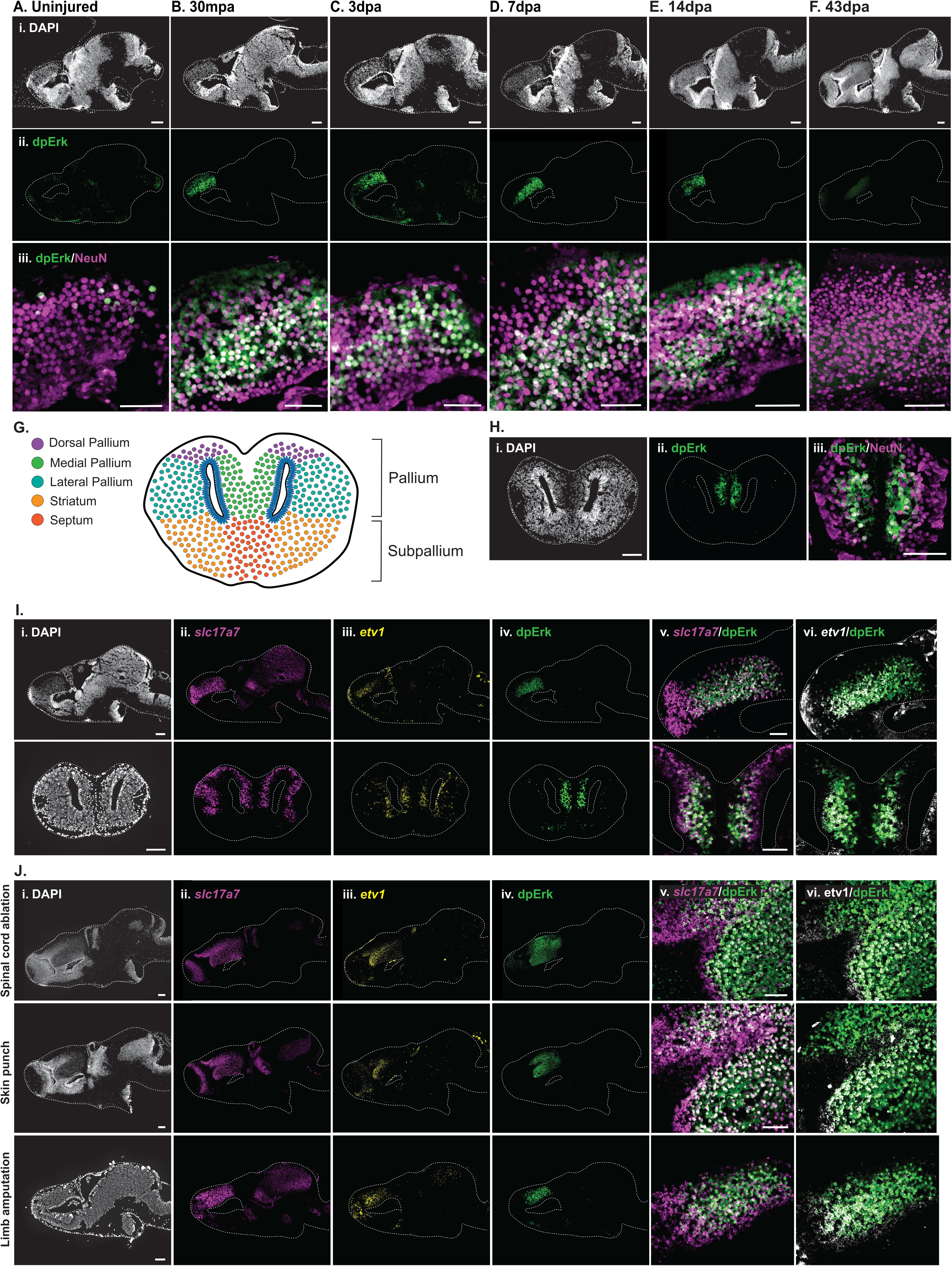
Injury leads to long-lasting Erk activation in the axolotl brain. **A-F)** Immunostaining using a dpErk antibody establishes the time-course for Erk activation in the brain at various time-points after a tail amputation. mpa = minutes post amputation. dpa = days post amputation. Scale bars: 200μm. Scale bars for iii images: 100μm. **G)** Schematic demonstrates the different regions of the axolotl telencephalon. The dorsal, medial and lateral regions make up the pallium, while the striatum and septum are found in the subpallium. **H)** Immunostaining for dpErk using a cross section of the axolotl brain at 3-days post tail amputation. Scale bar: 200μm. Scale bar in iii image: 100μm. **I)** *In situ* hybridization for both *slc17a7* and *etv1* paired with dpErk immunostaining using both longitudinal sections (top panel) and cross sections (bottom panel) of the axolotl brain. Scale bars: 200μm. Scale bars for v-vi images: 100μm. **J)** Erk activation in the brain is also demonstrated in *etv1*^+^/*slc17a7*^+^ neurons in response to differing injuries, including a spinal cord ablation, skin punch biopsy and limb amputation. Scale bars: 200μm. Scale bars for v-vi images: 100μm.

The axolotl brain can be roughly divided into five distinct regions that are more easily discernable in cross sections. These include the medial, dorsal and lateral pallium, along with the septum and striatum (**Fig. 1G**). To better establish the precise localization of dpErk signal within these brain regions, we performed dpErk immunostaining on cross sections at 3-days post tail amputation, and discovered that dpErk signal was restricted to the medial pallium of the axolotl brain (**Fig. 1H**).

While these immunostaining procedures documented a consistent increase in Erk signaling in response to a tail amputation in larval animals, we next wanted to determine whether a similar phenomenon was also found in adults. We performed a tail amputation on adult axolotls, and collected the brain tissues from these animals at an early timepoint of 1-hour post tail amputation, and at 3-days post tail amputation. In larval animals, increases in dpErk activity were consistently found at both timepoints (**Fig.S1B,D**). In adults, however, dpErk signal was not detected in the medial pallium at 1-hour post tail amputation, but was instead found in a few neurons in the dorsal pallium (**Fig.S1C**). At 3-days post tail amputation, we found abundant dpErk signal in the medial pallium of the telencephalon of adult animals (**Fig.S1E**). This demonstrates that neuronal activation in the brain in response to injury is a conserved phenomenon that is found in axolotls throughout their life but is activated more slowly in adults.

To better characterize the identity of these dpErk^+^ neurons, we performed fluorescent *in situ* hybridization to detect the expression of various marker genes. Recent single cell sequencing studies have demonstrated that the axolotl medial pallium is primarily composed of glutamatergic neurons ^58–60^. Moreover, as Erk signaling is typically associated with excitatory neurons, we designed a probe to detect vesicular glutamate transporter 1 (*slc17a7*) to identify glutamatergic neurons. While the identity of different glutamatergic subpopulations in the axolotl brain remains elusive, previous work has identified a few markers for long-range projection neurons that reside in the medial pallium of the axolotl brain ^61^. More specifically, *etv1*, a marker for long-range projection neurons that extend from the brain into the spinal cord in primates ^62^, is expressed in the medial pallium of adult axolotls ^61^. Thus, we performed a fluorescent *in situ* hybridization for *etv1* and *slc17a7* in conjunction with dpErk immunostaining. Using both longitudinal and cross sections to better asses the expression patterns of these genes, we discovered that dpErk^+^ neurons express both *etv1* and *slc17a7* (**Fig. 1I**), demonstrating that these cells of interest are indeed glutamatergic neurons that express mammalian markers for long-range projection neurons.

After better characterizing our neurons of interest, we next sought to understand the specificity of Erk activation in response to injury. To determine the specificity of this response, we performed different types of injuries that elicit a regenerative response in the axolotl, including a limb amputation, skin punch biopsy, or a spinal cord ablation, in which a small portion of the spinal cord is removed rather than the entire tail. After performing dpErk immunostaining on brain tissues isolated from animals with these different injuries, we consistently documented a robust increase in dpErk signaling within *etv1*^+^/*slc17a7*^+^ neurons in the medial pallium (**Fig. 1J, Fig.S2**). This demonstrates that neuronal Erk activation in the brain is not specific to a spinal cord injury, but rather represents a more global response to injury.

### Erk activation in the brain is necessary for normal tail regeneration

To better understand the functional importance of this neuronal activation in the brain in response to injury, we injected the axolotl brain with FR180204, a highly selective Erk inhibitor ^63^, or the corresponding vehicle control and then performed a tail amputation. After injecting the axolotl brain with FR180204 or the vehicle control, we measured tail regenerate lengths over a 21-day period. We discovered that inhibiting Erk signaling in the brain significantly reduced tail regenerate lengths, and that this difference became more pronounced over time (**Fig. 2A,B**). To more specifically explore the effect on axon regeneration, we performed β-III-tubulin staining on wholemount tails after injection with FR180204 or the vehicle control into the brain. The axons from both control and FR180204-injected animals appeared similar in their gross morphology, length and abundance at 7-days post tail amputation, and extended normally into the newly regenerated tail tissue (**Fig. 2C**). At 21-days post tail amputation, however, there appeared to be less axons re-growing into the regenerating tail in Erk inhibitor-injected animals compared to controls (**Fig. 2C**). This indicates that neuronal activation in the brain is necessary for axon regeneration in the tail, and that inhibiting this activation reduces the number of axons that extend into the newly regenerated tail tissue.

**Figure 2.**
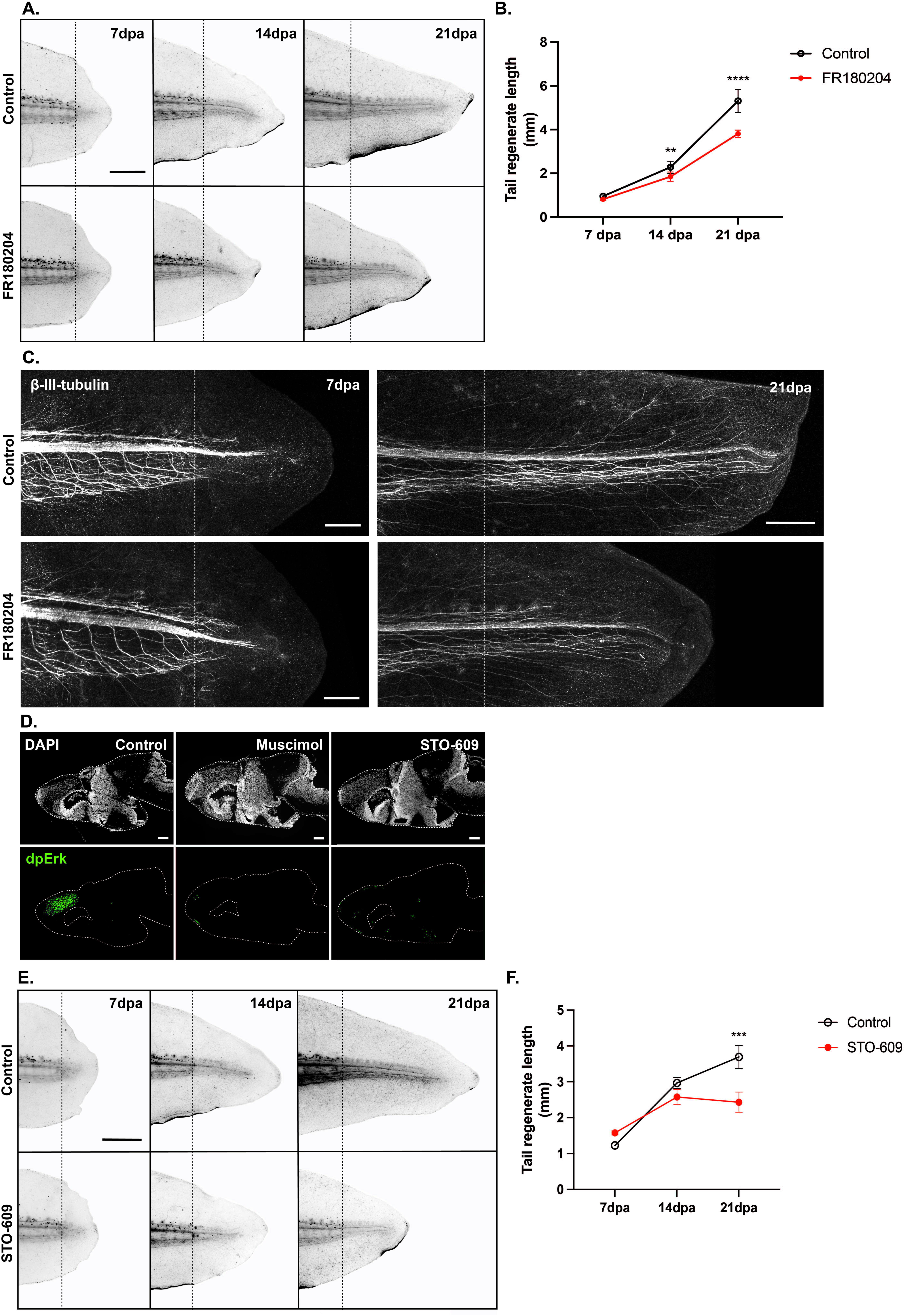
Erk activation in the brain is necessary for tail regeneration. **A)** Representative images of tail regeneration after injection with 10μm FR180204 or the corresponding vehicle control (0.1% DMSO) into the brain. Black dotted line indicates the original plane of amputation. Scale bar: 1mm. **B)** Graph demonstrates that injecting the axolotl brain with 10μm FR180204 significantly reduces tail regenerate lengths at 14dpa and 21dpa in comparison to vehicle controls (0.1% DMSO). *p<0.05. ****p<0.0001 (Two-Way ANOVA). **C)** β-III-tubulin antibody staining on regenerating wholemount tail tissues after injection with 10μm FR180204 or the vehicle control (0.1% DMS0) into the brain. White dotted line indicates the original plane of amputation. Scale bars: 500μm. **D)** Immunostaining with a dpErk antibody demonstrates that injection with 1mM Muscimol or 40μM STO-609 into the brain reduces Erk activation after a tail amputation. Scale bars: 200μm. **E)** Representative tail regenerate images after injection with 40μM STO-609 into the brain. Black dotted line indicates the original plane of amputation. Scale bar: 1mm. **F)** Graph demonstrates that injection with 40μM STO-609 into the brain significantly reduces tail regenerate lengths in comparison to vehicle controls (0.4% DMSO) at 21dpa. *** indicates p<0.001. (Two-Way ANOVA).

After identifying the importance of Erk signaling in the brain after injury, we next sought to determine the upstream regulators of Erk activation in the brain. As we identified these dpErk^+^ cells as excitatory glutamatergic neurons, we injected the axolotl brain with a highly selective GABA agonist, muscimol ^64^, to inhibit the excitability of glutamatergic neurons. After injection of muscimol into the medial pallium of the telencephalon, we performed a tail amputation to activate Erk signaling. Interestingly, we discovered that injection with muscimol completely abolished Erk activation in the brain, whereas vehicle controls still demonstrated a robust increase in Erk signal (**Fig. 2D**). This finding demonstrates that electrical activity is necessary for dpErk activation in the brain after injury.

In mammalian systems, calcium signaling has become a well-established upstream regulator of Erk signaling, acting through calmodulin-dependent kinases ^65,66^. To determine whether this signaling pathway is conserved in axolotls, we injected the axolotl brain with STO-609, a potent calmodulin-dependent kinase kinase (CaMKK) inhibitor ^67^, and then performed a tail amputation. We discovered that injection with a CaMKK inhibitor dramatically reduced Erk activation in the axolotl brain in comparison to vehicle controls (**Fig. 2D**). To determine whether this inhibition affected tail regeneration, we measured tail regenerate lengths over a 21-day period after injection with STO-609 in the brain. Similar to that of the Erk inhibitor, injection with STO-609 led to a reduction in tail regenerate lengths, which became more pronounced over time (**Fig. 2E,F**). Collectively, this work demonstrates that both electrical activity and CaMKK-regulated calcium signaling are required for Erk activation and act upstream of this signaling pathway.

### Neurons in the medial pallium extend axons to the hypothalamus

To better understand the identity and role of these dpErk^+^ neurons in regenerative repair, we used viral tracing to visualize the axons that emanate from medial pallium neurons. Recently, adenovirus has been shown to be an effective method of specifically labeling neurons in several amphibians ^68,69^. We injected adeno-associated viral particles with GFP under the control of a ubiquitous CAG promoter into the medial pallium of the axolotl brain, then used vibratome sections to image our labelled brain tissue. Using this technique, we discovered that neurons in the medial pallium projected axons in thick bundles down the anterior and posterior sides of the ventricle into the subpallium (**Fig. 3B**, yellow arrows; **Fig.S3**). While labelled neurons in the medial pallium predominantly extended axons into the ventral surface of the diencephalon, we also detected a few axons that extended into the brainstem, although these projections were sparse (**Fig. 3C**, yellow arrow; **Fig.S3**). To confirm that the viral labeled neurons do indeed upregulate dpErk after injury we performed antibody staining of animals containing GFP labeled neurons in the brain 1-day post tail amputation and verified that the GFP viral labelled axons in the pallium upregulated dpErk injury (**Fig. 3D**).

**Figure 3.**
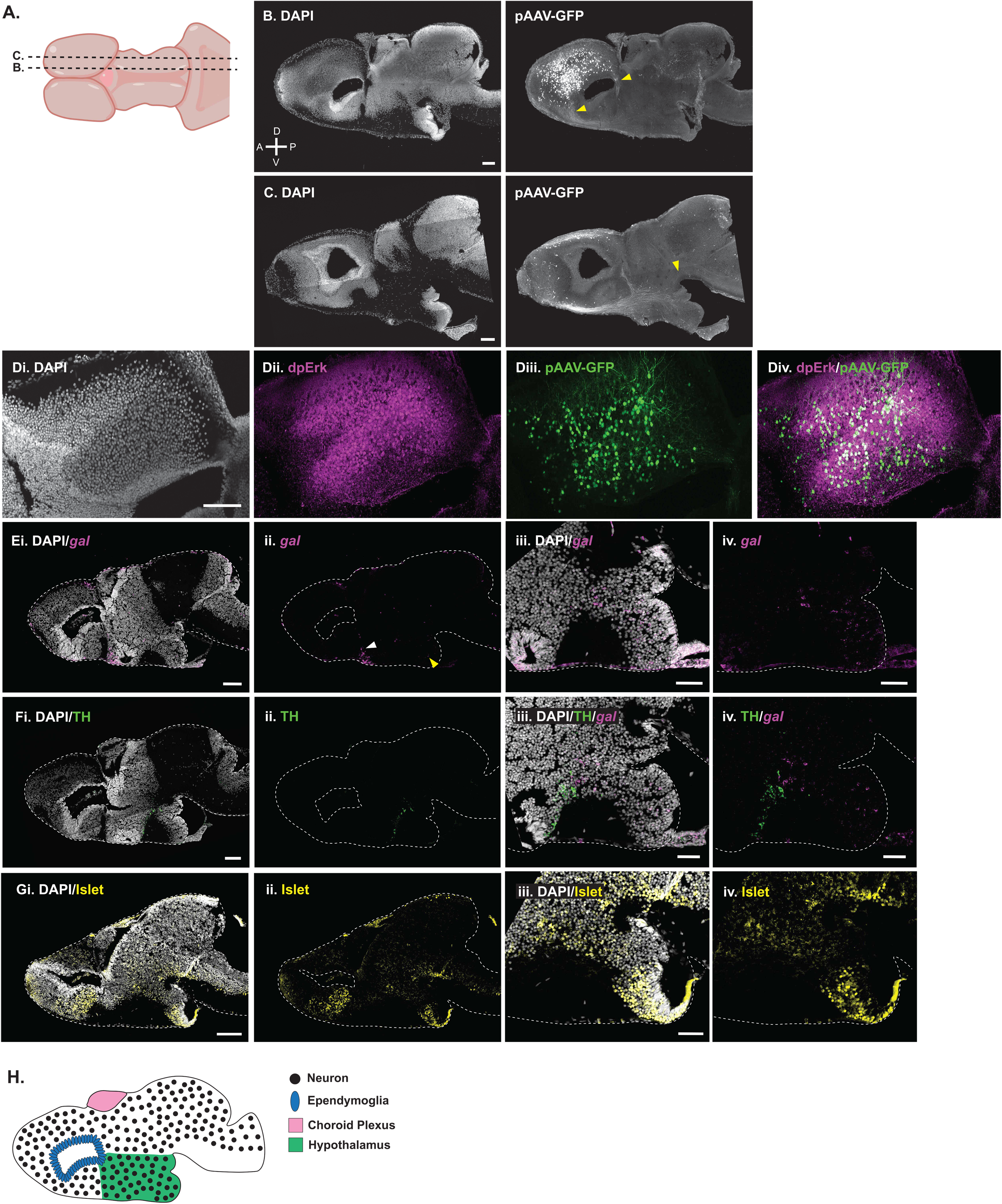
Neurons within the medial pallium extend axons into the hypothalamus. **A)** Schematic displays the approximate location of longitudinal vibratome sections that exhibit pAAV-GFP signal. **B, C)** Representative images of vibratome sections demonstrating the labelling of medial pallium neurons (B) and the axons that emanate from these neurons into the ventral surface of the subpallium (C). A few axons that extend into the spinal cord were also detected (C, yellow arrow). D= dorsal, V=ventral, A=anterior, P=posterior. Scale bars: 200μm. **D)** Immunostaining for dpErk in the brain of axolotls injected with pAAV-GFP at 1-day post tail amputation. Scale bar: 100μm**. E)** *In situ* hybridization for the hypothalamus marker, *gal*, demonstrates high *gal* expression immediately below the ventricle (ii, white arrow). At lower levels, *gal* is also detected in cells within the posterior portion of the subpallium (ii-iv, yellow arrow). **F)** Immunostaining for tyrosine hydroxylase (TH), a marker for dopaminergic neurons, demonstrates TH expression within the subpallium. Co-staining with TH and *gal* shows that both markers are localized in a similar region of the subpallium, but are not expressed in the same cell types (ii,iv). **G)** Immunostaining for Islet-1 demonstrates its broad expression in the subpallium. Scale bars: 200μm. Scale bars for iii-iv images: 100μm. **H)** Schematic displays the approximate location of the hypothalamus in the axolotl brain.

Recently, magnetic resonance imaging was utilized to generate a blueprint for the different regions of the axolotl brain ^70^. Through this line of work, the ventral surface of the diencephalon was identified as the hypothalamus due to its gross morphology and position, which appears similar to that of other amphibians. To further confirm that neurons in the medial pallium of the axolotl brain extend axons into the hypothalamus, we stained the axolotl brain for a few well-established markers that are expressed in hypothalamus-specific neurons in mammalian systems. We performed fluorescent *in situ* hybridization for galanin (*gal*), a neuropeptide that is abundant in the hypothalamus ^71–77^, in conjunction with a protein immunostain for tyrosine hydroxylase (TH), a marker for dopaminergic neurons. Moreover, we also performed immunostaining for Islet-1, which is abundant in subpallium neurons ^78^ and the hypothalamus ^79^. We discovered that *gal* was expressed in discrete populations of cells in the subpallium, ranging from neurons directly beneath the ventricle (**Fig. 3E**, white arrow) to the posterior side of the brain (**Fig. 3E**, yellow arrow). TH-expressing cells were found in a more compact, but similar region of the subpallium; indeed, dopaminergic neurons were also found in the posterior side of the brain, but were not expressed in *gal*-containing cells (**Fig. 3F)**. Islet-1 expression was detected in a majority of cells within the subpallium, and was particularly abundant within cells beneath the ventricle and the posterior region of the brain (**Fig. 3G**). While *gal* and TH appeared to mark distinct populations of neurons within the subpallium, their expression patterns along with Islet-1 allowed us to determine the general boundaries of the hypothalamus. Based on *gal,* TH and Islet-1 expression, the hypothalamus appears to extend along the subpallium from the ventricle to the posterior side of the brain (**Fig. 3H**).

### Transcriptome changes in etv1^+^/slc17a7^+^ neurons in response to injury

To uncover the downstream signaling pathways that are activated in dpErk^+^ neurons in response to injury, we utilized single cell sequencing to profile the transcriptome of these cells. We collected the telencephalon from larval axolotls using four different treatment groups, including 1) uninjured control animals, 2) 1-day post tail amputation, 3) 1-day post limb amputation, and 4) 1-day post tail amputation paired with muscimol injection into the brain to inhibit Erk signaling. Together, these four conditions aimed to uncover the signaling pathways that are upregulated in response to different injury contexts, and to identify those that are specifically regulated by Erk activation. We first annotated the broad cell types that exist in the axolotl telencephalon based on the expression of well-established marker genes that have been recently used to profile cells in the salamander brain ^58–60^. We identified ten distinct cell types, including ependymoglia (*gfap*^+^/*sox2*^+^), glutamatergic neurons (*slc17a7*^+^ or *slc17a6*^+^/*tubb3*^+^/gfap^-^), GABAergic neurons (*gad1*^+^/*gad2*^+^/*tubb3*^+^/gfap^-^), progenitor glutamatergic (*slc17a7*^+^ or *slc17a6*^+^/*tubb3*^+^/gfap^+^) or GABAergic (*gad1*^+^/*gad2*^+^/ *tubb3*^+^/gfap^+^) neurons, immune cells (*csf1r*^+^/*c1qb*^+^/*cd68*^+^ or *prf1*^+^/*cst7*^+^), oligodendrocytes (*olig2*^+^), oligodendrocyte progenitor cells (*cspg4*^+^/*pdgfra^+^*), endothelial cells (*cdh5*^+^/*cav1*^+^/*pecam1*^+^), and choroid plexus epithelial cells (*kcnj13*^+^)(**Fig. 4A,B**).

**Figure 4.**
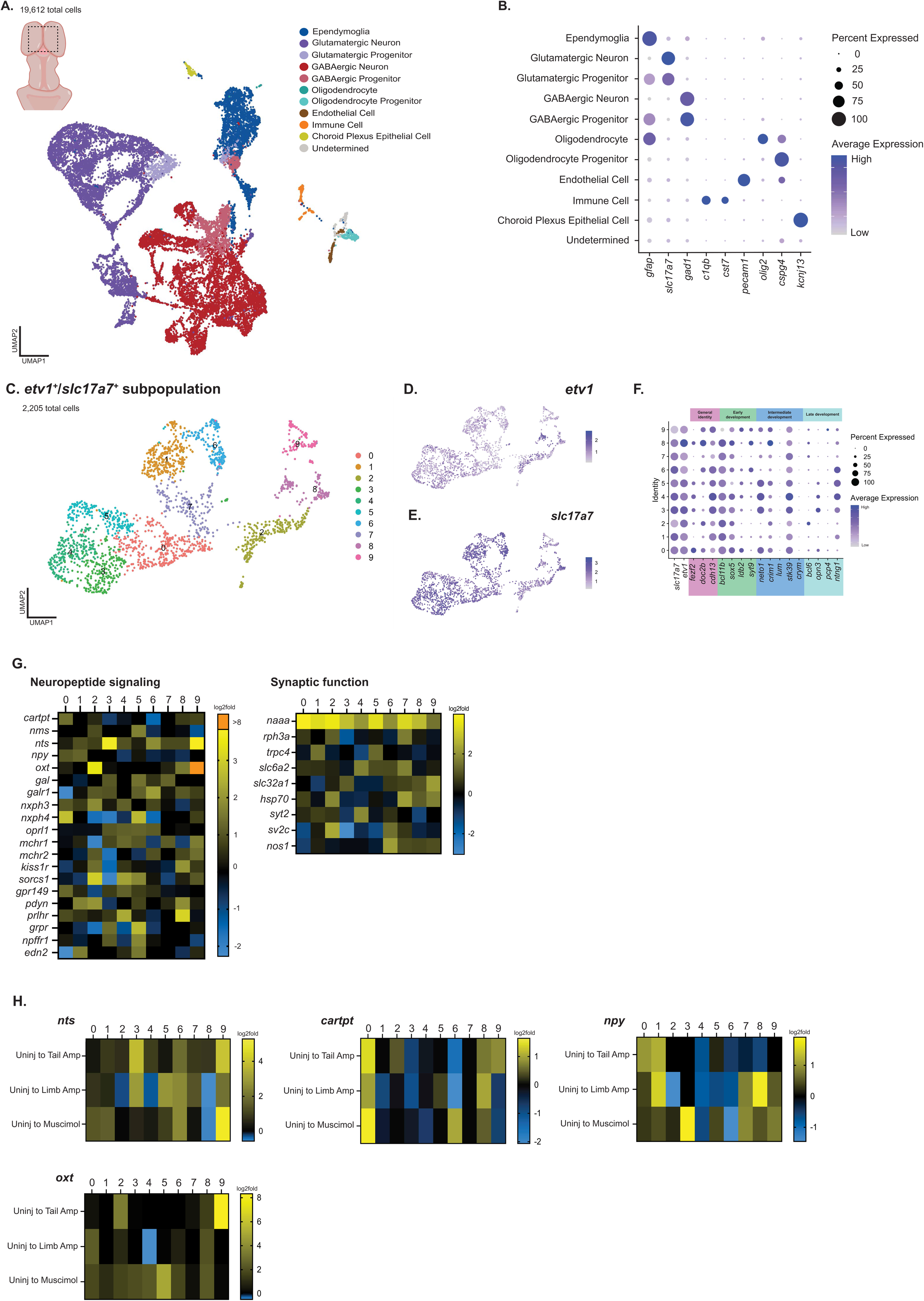
Single cell sequencing of the axolotl telencephalon identifies changes in gene expression after injury. **A)** Schematic highlighting the region of the axolotl telencephalon used for scRNA-seq. UMAP plot depicts the cell type composition of the axolotl telencephalon. **B)** Dot plot displays the expression of marker genes used to annotate cells in the axolotl brain. **C)** UMAP plot displays ten *etv1*^+^/*slc17a7*^+^ cell clusters. **D-E)** UMAP plots of the marker genes *slc17a7* and *etv1*. **F)** Dot plot displays the expression of mammalian spinal cord-projecting neuron markers within the ten *etv1*^+^/*slc17a7*^+^ axolotl cell clusters. **G)** Heatmaps display the log2fold change in expression of candidate genes involved in neuropeptide signaling or synaptic function between each cell cluster at 1-day post tail amputation compared to uninjured controls. **H)** Heatmaps depict log2fold changes in the expression of *nts, cartpt, npy* or *oxtl* between treatment groups.

We further subclustered ependymoglia (gfap^+^/sox2^+^) into 18 distinct clusters that were labelled as either active (notch1^+^/pcna^+^/mki67^+^/fgfr3^+^/wnt5b^+^) or quiescent (edn3^+^/lrtm4^+^) based on the expression of cell cycle genes or previously established markers (**Fig.S4A-C**) ^58^. As ependymoglia act as stem cells that can give rise to several cell types of the axolotl brain ^58^, we also subclustered these ependymoglia based on their cell fate. Some ependymoglia expressed markers for oligodendrocytes (olig2^+^/cspg4^+^), choroid plexus epithelial cells (kcnj13^+^), or GABAergic (gad1^+^) or glutamatergic (slc17a7^+^ or slc17a6^+^) neurons. Thus, we labelled these clusters as progenitor cells for these mature cell types (**Fig.S4D,E**). Next, we further subclustered mature glutamatergic (**Fig.S4F-H**) and GABAergic neurons (**Fig.S4I-K**). We identified 21 distinct glutamatergic subclusters, and 18 distinct GABAergic cell clusters. Each of these clusters were found across our treatment groups, with no obvious expansion of any cell type between treatments.

After annotating these broad cell types in the axolotl telencephalon, we subclustered *etv1*^+^/*slc17a7*^+^ neurons to better understand the transcriptomic changes that occur as a result of an injury in these cells. Our *in situ* hybridization experiments demonstrated that while *etv1* was predominantly expressed in glutamatergic neurons in the medial pallium, there was some additional, albeit minimal, expression in ependymal cells and GABAergic neurons (**Fig. 1G**). Thus, we subclustered *etv1*^+^/*slc17a7*^+^/*gfap*^-^/*gad1*^-^ cells to ensure only glutamatergic neurons were being assessed. This led to the generation of 10 distinct cell clusters (**Fig. 4C**) that expressed both *etv1* and *slc17a7* (**Fig. 4D, E**).

After clustering our cells of interest, we next wanted to investigate the expression of additional mammalian-specific long-range projection neuron markers. In salamanders, the pallium has been described as homologous to the mammalian cortex ^80^, while the medial pallium has been more specifically identified as homologous to the mammalian hippocampus ^81^. Recently, single cell profiling of the axolotl telencephalon described different populations of glutamatergic neurons in the medial pallium that closely resembled both hippocampal- and cortex-derived neurons ^58^.

In mammals, neurons born within the cortex can extend axons within the cortex, or to the spinal cord or thalamus. More specifically, mammalian etv1^+^ neurons within the cortex have been shown to extend into either the spinal cord or thalamus, while we discovered that axolotl etv1^+^ neurons in the medial pallium primarily extend axons into the hypothalamus. Collectively, this suggests that *etv1* is not a conserved marker for spinal-cord projecting neurons in the axolotl. To better address whether axolotl etv1^+^ cells express other mammalian markers for long-range projection neurons, we looked at the expression of markers for cortex-born neurons that extend to the spinal cord. We discovered that axolotl *etv1*^+^/*slc17a7*^+^ neurons express multiple mammalian marker genes for spinal-cord projecting neurons that are found in mice at various stages of development, including *cdh13*, *bcl11b*, *sox5*, *neto1*, *stk39* and *ntng1* (**Fig. 4F**). As cortex-derived neurons in mammals can also project within the cortex or extend to the thalamus, we also examined the expression of markers for cortex- and thalamus-projecting neurons (**Fig.S5A**). We discovered that axolotl *etv1*^+^/*slc17a7*^+^ neurons also express a number of thalamus-projecting neuron markers, but largely lack cortex-projecting markers. Collectively, this indicates that many mammalian long-range projection neuron markers may not be conserved in the axolotl.

To determine whether axolotl *etv1*^+^/*slc17a7*^+^ neurons instead express mammalian markers for hippocampal-derived neurons, we again examined the expression of various hippocampus marker genes. Indeed, we found that axolotl *etv1*^+^/*slc17a7*^+^ neurons appear to express multiple general-identity hippocampal marker genes, including *bdnf*, *cabin1*, *gria1*, *gria2*, *gria3*, and *grm5* (**Fig.S5B**).

After investigating the conservation of various mammalian marker genes, we next examined the transcriptome changes that occur in the brain as a result of a tail amputation. We first identified differentially expressed genes in each *etv1*^+^/*slc17a7*^+^ cell cluster at 1-day post tail amputation in comparison to uninjured controls. We performed a GOrilla gene ontology analysis on these differentially expressed genes and identified multiple biological processes that are activated after tail amputation in multiple cell clusters. These biological processes included neuropeptide signaling (**Fig. 4G**), and processes involved in synaptic function such as neurotransmitter loading into the synapse and signal release from synapse (**Fig. 4G**).

As neuropeptides can often act as neurotransmitters in the brain to activate synaptic terminals ^82–87^, and many have been shown to activate hormone-producing neurons in the hypothalamus ^88–92^, we decided to further explore the expression patterns of multiple neuropeptides, including neurotension (*nts*), neuropeptide y (*npy*), oxytocin (oxt), and cocaine- and amphetamine-regulated transcript (*cartpt*). Each of these candidate neuropeptides exhibited large log2fold increases in their expression in at least one *etv1*^+^/*slc17aa7*^+^ cell cluster at 1-day post tail amputation, indicating that their expression may be regulated by Erk signaling. To further explore their expression patterns across our different treatment groups, we compared the log2fold changes in their expression at 1-day post tail amputation, 1-day post limb amputation, and 1-day post tail amputation after injection with muscimol, all compared to uninjured controls (**Fig. 4H**). While *nts* appeared to be upregulated in a majority of these cell clusters after tail amputation, some cell clusters also exhibited a similar trend after limb amputation. For example, in cell cluster 3, *nts* exhibited a >3 log2fold increase after tail amputation, and a >2 log2fold increase after limb amputation; while no change in expression was shown in muscimol-injected tissues. This suggests that increases in *nts* expression appear as a result of Erk activation in this specific cell cluster, regardless of the injury context. In cell cluster 5, a >2 log2fold increase in *nts* expression was only demonstrated after limb amputation, indicating that this change in expression may in fact be specific to limb amputation (**Fig. 4H**). Collectively, these results demonstrate that increases in *nts* expression in response to tail or limb amputation vary between different subpopulations of *etv1*^+^/*slc17a7*^+^ neurons.

We next assessed the expression of other neuropeptides between our different treatment groups, including *cartpt*, *npy*, and *oxt*. Both *cartpt* and *npy* appeared to be upregulated in a few cell clusters in response to tail amputation, but upon further analyses, similar trends were also shown in the same cell clusters in response to limb amputation and muscimol injection (**Fig. 4H**). Ultimately, this indicates that both neuropeptides may not be regulated by Erk signaling. In *oxt* comparisons, *oxt* exhibited a dramatic upregulation in cell clusters 2 and 9 that was specific to tail amputation, however, there were no significant changes in *oxt* expression in the remaining *etv1*^+^/*slc17a7*^+^ cell populations (**Fig. 4H**). Given that *nts* was upregulated in a majority of *etv1*^+^/*slc17a7*^+^ cell populations after a tail amputation, we decided to further pursue the role of this neuropeptide in axolotl tail regeneration.

After uncovering interesting *nts* expression dynamics between *etv1*^+^/*slc17a7*^+^ cell clusters, we next sought to further validate the expression of this neuropeptide in the axolotl brain. We performed fluorescent *in situ* hybridization for *nts* after a tail or a limb amputation in conjunction with *gal* to mark the hypothalamus **(Fig. 5A,B**). We discovered that *nts* was localized in the subpallium, and medial and lateral pallium (**Fig. 5A**). After both tail and limb amputation, we also discovered *nts* was expressed in low levels in axons within the medial pallium (**Fig. 5A,Bvi** yellow arrow), and was abundant within the cell bodies of a subpopulation of neurons in the medial pallium that also expressed *etv1* (**Fig. 5A,Bvi**; white arrow). This indicates that both limb and tail amputation lead to the production of *nts* in a subpopulation of neurons within the medial pallium.

**Figure 5.**
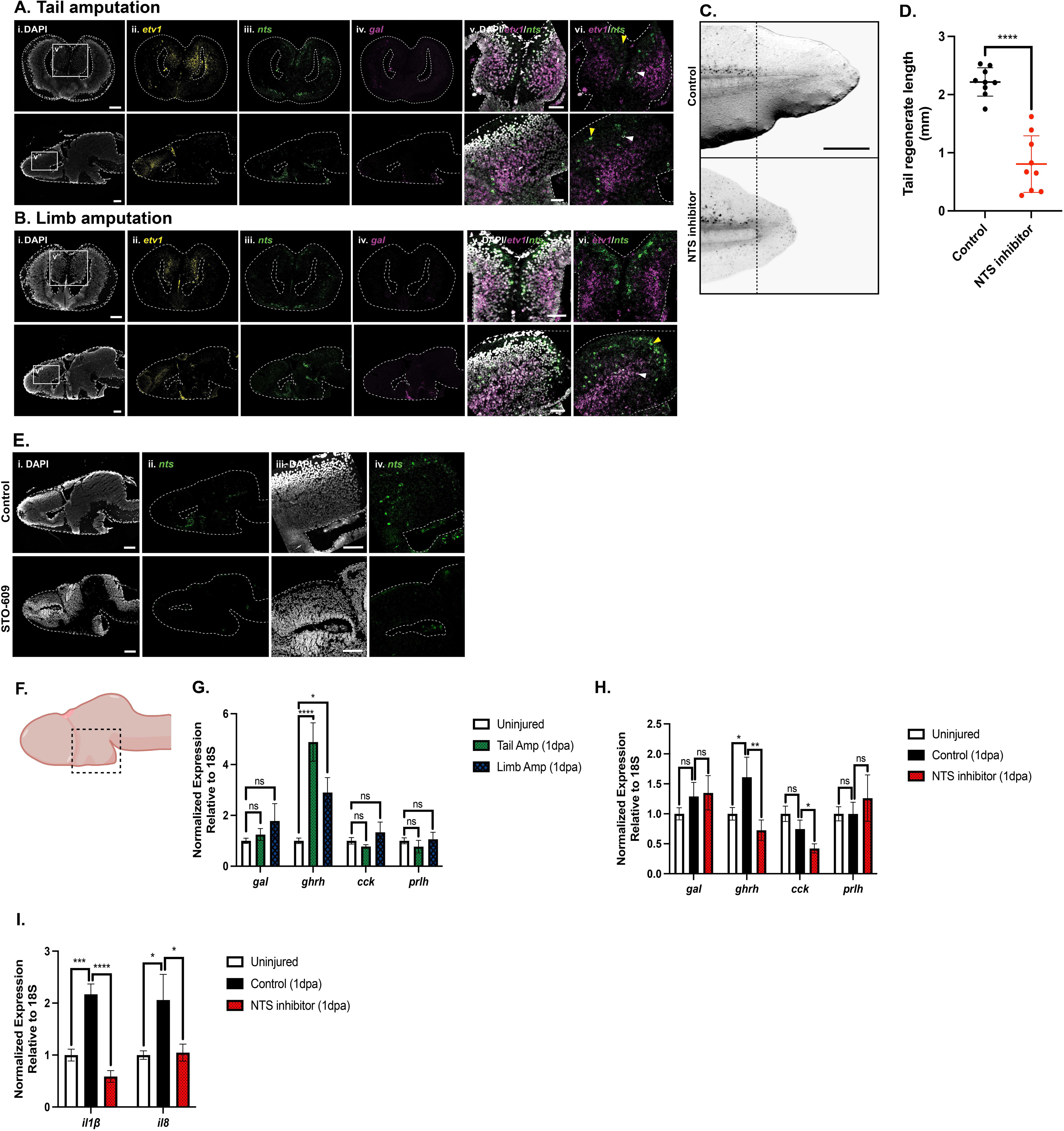
Neurotensin is essential for axolotl tail regeneration. **A,B)** *In situ* hybridization using both longitudinal and cross sections after tail (A) or limb (B) amputation demonstrates the expression patterns of *etv1*, *nts* and *gal* in the axolotl brain. Scale bars = 200μm. Scale bars for v, vi images: 100μm. **C)** Representative images of control and neurotensin inhibitor-treated animal tail regenerates at 12-days post tail amputation. Scale bar: 1mm. **D)** Graph demonstrates that inhibiting neurotensin significantly reduces tail regenerate lengths at 12-days post tail amputation. ****p<0.0001 (t-test). **E)** *In situ* hybridization displays a reduction in *nts* expression in the axolotl brain after injection with 40μM STO-609 in comparison to vehicle controls. **F)** Schematic indicates the approximate location of hypothalamus tissue isolated for qPCR experiments in panels G and H. **G)** qPCR demonstrates the expression of multiples hormones within the hypothalamus in uninjured controls and at 1-day post tail and limb amputation. ns = not significant. *p<0.05. ****p<0.0001 (One-Way ANOVA). **H)** qPCR displays the expression of multiple hormones within the hypothalamus of uninjured controls and at 1-day post tail amputation after treatment with the neurotensin inhibitor or the corresponding vehicle control. ns = not significant. *p<0.05. **p<0.01. (One-Way ANOVA). **I)** qPCR demonstrates the expression of inflammatory response genes in the tail tissue of uninjured animals and at 1-day post tail amputation in neurotensin inhibitor-treated animals or vehicle controls. *p<0.05. ***p<0.001. ****p<0.0001. (One-Way ANOVA)

### Neurotensin upregulation is necessary for successful tail regeneration

To further examine the role of neurotensin during tail regeneration we inhibited neurotensin using the well-characterized neurotensin receptor 1 antagonist SR142948A ^93–95^. Before amputation, the antagonist was microinjected into the brain, and we then followed the animals for 12 days post-amputation and quantified the length of tail regenerates. When neurotensin was inhibited we found that the tail regenerates from these animals were significantly shorter than controls (**Fig. 5C,D**). This phenotype indicates that neurotensin is essential in the early phases of regeneration; to establish where in the pro-regenerative molecular circuitry neurotensin lies we carried out fluorescent *in situ* hybridization for *nts* after Erk inhibition. Interestingly, we found that *nts* mRNA was not up-regulated in the medial pallium or subpallium after tail amputation in comparison to control animals (**Fig. 5E**). This data indicates that an upregulation of dpErk after injury is necessary to induce activation of *nts* transcription.

To gain a better understanding of the role of neurotensin in regenerative repair, we quantified the levels of multiple hormones that are produced in the hypothalamus that have been documented in other model systems to act downstream of neurotensin ^96–99^. To examine the changes in hormone production after injury, we performed qPCR on microdissected hypothalamus tissue from uninjured controls and at 1-day post tail and limb amputation (**Fig. 5F**). Interestingly, we found only growth hormone releasing hormone (*ghrh*) was significantly upregulated in the hypothalamus in response to injury (**Fig. 5G**), while cholecystokinin (*cck*), galanin (*gal*) and prolactin-releasing hormone (*prlh*) were unaffected. We next looked at the levels of the same hormones after neurotensin inhibition and discovered that inhibiting neurotensin significantly reduced *ghrh* expression, indicating that *ghrh* production in the hypothalamus is regulated by neurotensin (**Fig. 5H**). We also found that *cck* was significantly downregulated in response to neurotensin inhibition, indicating a possible interaction between *cck* and neurotensin (**Fig. 5H**). Together, these results suggest that upregulation of neurotensin is necessary to activate the production of hormones within the hypothalamus.

In mammals, neurotensin has a well-documented role in the regulation of the immune response ^100–106^, which is known to play an important role in salamander regeneration ^107–111^. To determine whether neurotensin may be regulating the axolotl immune response after injury, we looked at two key regulators of the inflammatory response, including interleukin 1β (*il1*β) and interleukin 8 (*il8*) in the axolotl tail after a tail amputation. Both genes were significantly upregulated at 1-day post tail amputation, however, when neurotensin was inhibited, levels of both genes remained at homeostatic levels (**Fig. 5I**). Together, this suggests both *il1*β and *il8* are downstream of neurotensin in the regeneration program that must be executed for faithful tail regeneration to occur.

## Discussion

Much of the work on our understanding of the process of regeneration has come from studies focused on changes that occur in cells adjacent to the wound site. Here we have focused on changes that occur in cells in the brain in response to a distal injury. We identified the dual phosphorylation of Erk as a key change that occurs in a subpopulation of neurons in the brain after injury. To identify the specific neuronal populations in which Erk is necessary to drive regeneration, we carried out single cell sequencing on isolated brain tissue in a variety of conditions and later used *in vivo* modulation of gene function to identify downstream drivers of regeneration.

A central finding of our work highlights a complex brain-body axis of signaling that is necessary to execute a distal pro-regenerative response. We identified a population of dpErk glutamatergic neurons that synapse in the hypothalamus that drive the upregulation of several neuropeptides including neurotensin in response to a distal injury. These data are an invaluable resource to gain deeper insights into how the brain reacts, interprets and executes a pro-regenerative program. It has been well-established that nerves play a crucial role in the regenerative response, especially in the limb where several factors including newt anterior grade protein (NAG) and neuregulin (NRG1) are known to be secreted by nerves and stimulate proliferation in the limb blastema ^112–115^. Our work suggests that injury activates a signaling cascade in glutamatergic neurons in the medial pallium, which in turn activates many neuropeptides including neurotensin. Neurotensin, acting as a neurotransmitter, regulates many downstream factors which may be transmitted via axons or may activate the production of hormones, such as *ghrh*, that is transmitted via the blood stream to activate cells adjacent to the injury site. Work in other species including Xenopus and zebrafish has previously identified circulating hormones as playing a role in regeneration ^116–131^. In Xenopus, melanocortin 4 receptor (Mc4R) signaling is upregulated after limb injury in the regeneration blastema, and this is attributed to α-melanocyte-stimulating hormone (αMSH) production in the hypothalamus. Interestingly, in deinnervated limbs, administration of αMSH suffices to partially rescue blastema formation, suggesting an important early role for αMSH hormone production in regeneration ^132^. Additionally, Mc4R knock-out mice fail to regenerate mouse digit tips, suggesting this pathway is conserved and necessary for vertebrate regeneration ^133^. Circulating hormones like androgens and increasing levels of thyroid hormone are also known to affect the regenerative capacity of fins in some fish and in antler regeneration in deer ^118,122–131,134^. More recently, adrenergic signaling has been found to stimulate cell division in uninjured tissues in response to injury of one of the salamander’s other limbs ^48^. How factors like hormones that are produced in a very distant part of the body modulate signaling at an injury site is largely unknown. Work from planaria proposes that Erk signaling facilitates whole body regeneration, and that waves of Erk are transmitted via muscle in the animal’s body wall ^135,136^. Interestingly, in zebrafish scale regeneration Erk signaling has also been found to play a key role; however in this case they found waves of Erk travel in concentric rings across the regenerating scales ^137–139^ . Fascinating work in mammals has found Erk signaling also plays a key role in response to injury. In comparative work, prolonged Erk signaling was found to result in scar-free wound healing in the skin of the regeneration competent African spiny mouse, while in *Mus musculus* although they activated Erk after injury it rapidly declined and the mouse formed scar tissue ^140^. Importantly, when Erk signaling is impaired in the spiny mouse, the animal shifts from scar-free wound healing to fibrotic scarring ^140^.

Our work has uncovered a prolonged Erk response in a subpopulation of glutamatergic neurons in medial pallium which in the case of tail regeneration lasts up to ∼40 days post-injury. Impairing Erk signaling leads to defects in tail regeneration. Our work establishes a connection between how the brain responds to distal injury and identifies interesting differences in neurohormones activated when the limb is amputated versus tail amputation and molecules activated in response to both types of injury. In the future it will be important to decipher the circuitry which is activated to tell the brain to regenerate a limb versus a tail. Work in zebrafish and mice suggests that Fgf and ErB-2 are upstream of Erk and inhibition of these pathways leads to reduced Erk ^137,140^. In previous work investigating spinal cord regeneration in our lab we have identified Erk signaling to be activated in the glial cells adjacent to the injury site and found that changes in membrane potential were crucial to activate Erk ^57^. Here we have found that an influx of calcium is essential to activate Erk in the brain, however, where this calcium comes from is unknown. It will be important in the future to determine how the signal from the injury site is transported directly to the brain, and to determine whether this a mechanical signal or a rapid transcriptional response at the injury site that is transported via axons or the circulatory system to the brain. An interesting study on leg regeneration in cockroaches, whereby they severed the connections between the brain and the injured limb showed that this impaired limb regeneration and eliminated the delay in molting associated with regeneration, suggesting that input from the brain was necessary for successful limb regeneration ^141^. Work on many different regenerative organs has suggested that regeneration does not occur simply by activation of local cells, but that inter-organ communication is a shared feature of regeneration, reviewed in ^142^. It is still unknown how this inter-organ and long distance communication occurs, factors may be released from injured cells and carried in the blood stream, bioelectrical signaling has recently been shown to play an important role in development and regeneration as have factors released from nerves ^113,130,143–145^. In this paper we show that different types of injury lead to the activation of Erk signaling in a specific population of glutamatergic neurons in the pallium and after tail amputation these neurons activate neurotensin which is necessary for successful axon regeneration. In summary, this work reveals a complex interplay of distant and local responses to injury, highlighting the need to further study how the brain responses to distant injuries and to ultimately elucidate how this long-distance interorgan communication occurs.

## Methods

### Animal Care and Husbandry

All animals were bred and obtained at the Marine Biological Laboratory in accordance with IACUAC regulations. For all studies, the white strain of axolotls was utilized. Although the sex of larval animals cannot be determined, in experiments using adult animals, both female and male axolotls were utilized to account for potential sex-specific differences.

Prior to all experiments, larval animals (2-3cm) were anesthetized in 0.01% p-amino benzocaine (Sigma), while adult animals were anesthetized in 0.03% p-amino benzocaine. All amputations were performed using a number 10 scalpel; ∼3mm of tail tissue was amputated from larval animals, while ∼1 inch of tail tissue was amputated from adults. Limb tissue was amputated through the forearm approximately 1mm below the elbow. Skin biopsy punches were performed using a sterile 2 or 4-mm disposable biopsy punch with plunger (Miltex, York, PA, USA). Spinal cord ablation injuries were performed as previously described ^31,32,57^. After injury, animals were placed in individual cups and monitored for the duration of the experiments.

### Immunohistochemistry

Axolotl brain tissues were micro-dissected and fixed in 4% paraformaldehyde at 4°C overnight. The next day, tissues were washed 3x5min in phosphate buffered saline (PBS), then incubated in 15% sucrose for 10minutes. Samples were next incubated in 30% sucrose at 4°C overnight, then embedded in O.C.T. for cryosectioning. Cryosections were sliced at a 20μm thickness on a Leica 1850 Cryomicrotome and collected on Superfrost Plus microscope slides (Fisher).

For immunostaining procedures, slides were initially incubated in hot PBS at 70°C for 20min, then washed 2x5min in PBS at room temperature. Slides were next incubated in boiling citrate buffer for 10min, then permeabilized in 0.1% Triton/PBS for 30min. Following permeabilization, sections were blocked in blocking buffer (2% BSA, 2% goat serum in 0.1% Triton/PBS) for 1-hour, then incubated in primary antibody at 4°C overnight. Primary antibodies used in immunostaining procedures included an anti-mouse NeuN (1:100, Chemicon #MAB377) paired with an anti-rabbit dpErk (1:200, Cell Signaling, #A11000). The next day, slides were briefly washed 4x5min in 0.1% Tween-20/PBS (PBST), then incubated in secondary antibody (1:200) for 3 hours at room temperature. Secondary antibodies included AlexaFluor 488 goat anti-rabbit IgG (Invitrogen #A11008) and AlexaFluor 555 goat anti-mouse IgG (Invitrogen #A32727). After incubation in secondary antibodies, slides were washed 3x10min in PBST, counterstained with DAPI and mounted for imaging. All slides were imaged using a Zeiss 780 confocal microscope.

### Wholemount Immunostaining

Tail tissues were fixed in 4% paraformaldehyde at 4°C overnight, then washed 3x5min in PBS the following day. Tissues were next dehydrated in a graded series of methanol washes (25%, 50%, 75%) for 5min each, then stored in 100% methanol at −20°C until required.

For wholemount immunostaining, tails were rehydrated using a graded series of methanol washes (75%, 50%, 25%) and were then washed 3x5min in PBST. Tail tissues were next incubated in 2μg of proteinase K for 10 min at room temperature, and then further permeabilized in 0.2% Triton/PBS for 30min. Following permeabilization, tissues were blocked for 1hour in blocking buffer (2% bovine serum albumin, 10% goat serum in 0.2% Triton/PBS), and incubated in a β-III-tubulin primary antibody (1:500, Sigma #T8578) at 4°C overnight. The next day, tissues were washed thoroughly 3x30min in PBST and incubated in AlexaFluor 555 goat anti-mouse IgG secondary antibody (1:200, Invitrogen #A32727) at 4°C overnight. The following day, tails were again washed 3x30 min in PBST, then counterstained with DAPI. Finally, tails were embedded in 2% agarose and imaged on a Zeiss 780 confocal microscope.

### RNA Scope In Situ Hybridization

RNAscope *in situ* hybridization experiments were performed according to the manufacturer’s instructions (Advanced Cell Diagnostics). Cryosections were initially washed for 10min in PBS, and then baked for 30min at 60°C. Slides were next post-fixed in 4% paraformaldehyde for 15min at 4°C, then dehydrated in a graded series of ethanol dilutions (50%, 75%, 100%) for 5min each. Following dehydration, slides were air dried for 5min, then incubated in hydrogen peroxide for 10min at room temperature. Samples were briefly rinsed in deionized water, then incubated in hot target retrieval buffer at 85°C for 5min. After target retrieval, slides were rinsed in deionized water for 15s, washed in 100% ethanol for 3 min, and air dried for 5min at room temperature. Samples were next permeabilized using protease III for 30min at 40°C, briefly washed in deionized water, then hybridized with probes for 2 hours at 40°C. All probes were designed and distributed by Advanced Cell Diagnostics based on axolotl-specific mRNA sequences. Probe incubation included *etv1*-C1 paired with *slc17a7*-C2, or *etv1*-C1 paired with *nts*-C2 and *gal*-C3. After hybridization, slides were washed briefly in wash buffer, then incubated in 5x saline sodium citrate overnight at room temperature. The next day, slides were incubated at 40°C in Amp1 for 30min, Amp2 for 30min, and Amp3 for 15min. Samples were next incubated in HRP-C1 at 40°C for 15min, then treated with Opal-520 dye (1:750, Akoya #OP-001001) at 40°C for 30min to visualize *etv1*-C1. Next, slides were blocked with HRP blocker for 15min at 40°C, then incubated in HRP-C2 for 40°C for 15min. This was followed by an incubation in Opal-570 dye (1:750, Akoya #OP-001003) at 40°C for 30min to detect C2 probes. After blocking with HRP blocker for 15min at 40°C, slides were incubated in HRP-C3 for 40°C for 15min, then Opal-690 dye (1:100, Akoya #FP1497A) at 40°C for 30min to detect the C3 probe. Following a final blocking step in HRP blocker for 15min at 40°C, slides were counterstained using DAPI and mounted for imaging.

To detect dpErk protein expression in conjugation with *etv1* and *slc17a7*, the final HRP-C3 and Opal-690 dye incubation were omitted. Instead, after incubation in HRP blocker, slides were washed twice for 5min each with Tris-buffered saline (TBS). Samples were next blocked in blocking buffer (2% BSA/10% goat serum/0.1% Tween-20/TBS) for 1hour at room temperature, then incubated in dpErk antibody (1:200, Cell Signaling #A11000) at 4°C overnight. The following day, slides were washed in TBS, then incubated in AlexaFluor 647 goat anti-rabbit secondary antibody (1:200, Invitrogen #A21244) for 3hours at room temperature. After incubation in secondary antibody, slides were washed in TBS, counterstained using DAPI and mounted for imaging. All sections were imaged using a Zeiss 780 confocal microscope.

### Brain Injections and Viral Tracing

Prior to injection, AAV-CAG-GFP (AAV9)(Addgene, #37825-AAV9) viral particles were diluted 1:10 in 5μg/mL polybrene to facilitate viral uptake into animals, and pharmacological inhibitors were diluted to the appropriate working concentrations using PBS. Pharmacological inhibitors included 10μM FR180204, 40μM STO-609, 1mMol Muscimol, and the corresponding vehicles controls. A 1% Fast Green solution was also added to the viral particle or inhibitor-containing solutions to aid in injection visualization. These solutions were injected into the axolotl medial pallium using a PV 820 Pneumatic PicoPump pressure injector. After injection, tail amputations were performed as required. For animals injected with viral particles, axolotls were screened for successful GFP signal after 3 weeks.

For animals injected with the neurotensin inhibitor, axolotls were injected with 1 µM SR142948A into the ventricle of the brain before injury. The tail was then amputated and the animals were incubated in axolotl water containing the neurotensin inhibitor.

### Vibratome Sectioning and Staining

Brain tissues with positive AAV-GFP signal were micro-dissected and fixed in 4% paraformaldehyde at 4°C overnight. The next day, tissues were washed 3x5min in PBS, then embedded in 2% agarose. Brain tissues were sectioned at 80 μm using a Leica 1000S Vibratome. Free-floating vibratome sections were immediately placed in PBS within 24-well tissue culture plates (Fisherbrand) and processed for immunohistochemistry staining procedures.

Briefly, vibratome sections were washed 3x5min in PBS, then permeabilized with 0.1% Triton/PBS for 30min. Following permeabilization, sections were blocked for 1hour in blocking buffer (2% BSA, 2% goat serum in 0.1% Triton/PBS) and incubated in primary antibodies at 4°C overnight. Primary antibodies included an anti-mouse dpErk (1:200, Cell Signaling #5726S) and an anti-rabbit GFP (1:200, Invitrogen, #A11100). The next day, sections were washed 4x5min in PBST, then incubated in secondary antibodies for 3hours at room temperature. Secondary antibodies included an AlexaFluor 555 goat anti-mouse IgG (1:200, Invitrogen #A32727) and an AlexaFluor 488 goat anti-rabbit IgG (1:200, Invitrogen #A11008). Tissue sections were next washed 4x10min in PBST, counterstained with DAPI, and mounted on Superfrost Plus Microscope slides. Vibratome sections were imaged using a Zeiss 780 confocal microscope.

### Sample Preparation for Single Cell Sequencing

The telencephalon was micro-dissected from four larval animals for each treatment group and was roughly trimmed to exclude the olfactory bulb and the left/right sides of the telencephalon. Treatment groups included brain samples from uninjured animals, animals at 1-day post tail amputation and 1-day post limb amputation. An additional treatment group included brain tissue microdissected at 1-day post tail amputation that had been injected twice with 1mMol Muscimol. Muscimol injections into the medial pallium were performed at the time of tail amputation and the following day at 1 hour prior to microdissection.

After microdissection, a single-cell suspension was generated using the Whole Skin Dissociation Kit (Miltenyi Biotec) according to manufacturer’s instructions. Briefly, brain tissue was incubated in cell culture media containing 10% fetal bovine serum at room temperature until all microdissections were complete. After isolation, brain tissues were transferred into Buffer L/Enzyme P solution and gently mixed. Enzyme D and Enzyme A were then added to the solution and carefully mixed. Brain tissues were dissociated by further mixing on a GeneMate GyroMixer for 20min at room temperature. Throughout this 20min incubation, samples were occasionally pipetted using a p1000 to mechanically break apart any remaining tissue clumps. After 20min, the cell suspension was strained through a 70 μm cell strainer to remove any remaining tissue clumps. The single cell suspension was finally centrifuged at 300xg for 10min at room temperature, and the pelleted cells were resuspended in cell culture media containing 10% fetal bovine serum.

The Chromium Next GEM Single Cell 3’ Reagent Kit (Dual Index; 10X Genomics) was utilized to profile cells from the axolotl brain using a Chromium X (10x Genomics) instrument. All single cell GEM generation, barcoding, cleanup, cDNA amplification and library construction experiments were performed according to manufacturer’s instructions. LC Sciences carried out quality control and sequencing of the libraries, 400-500 M PE150 reads per sample (approx. 120Gbases/library). Reads were then aligned to the axolotl genome (AmexG.v6-DD) and assigned to genes based on annotation AmbMexT_V47 mapped to AmbMex60DD using Cell Ranger.

### Seurat Clustering and Analyses

For single cell analysis, 19,612 total cells were clustered and analyzed using the RStudio Seurat 5.1.0. The raw data from each of the 4-treatment groups were merged using the merge function to form 4 distinct layers. The layers were next normalized, and a linear transformation was applied to scale the data. A Principal Component Analysis (PCA) was next performed, and the top 20 PCs were selected based on an elbow plot of variance. These top 20 PCs were then used to generate a Uniform Manifold Approximation and Projection (UMAP) plot for visualization. Clustering was performed using the Louvain algorithm at a resolution of 6. Final cell clusters were annotated based on the expression of well-established marker genes ^58,59^. To run differential gene expression between samples, the 4 layers were integrated using the IntegrateLayers function. A subset of *etv1*^+^/*slc17a7*^+^ positive cells was generated using the subset function, in which *etv1* and *slc17a7* expression was set >0 while *gfap* and *gad1* expression was set =0. Ten cell clusters were generated for this *etv1*^+^/*slc17a7*^+^/*gfap*^-^/*gad1*^-^ subset using a resolution of 0.8, with a total of 2,205 cells. Ependymal glial cells were also subclustered using the subset function, with *sox2* and *gfap* expression set >0. This generated 18 cell clusters at a resolution of 0.8 with a total of 4,744 cells. Ependymal glial cells were annotated based on the expression of previously established marker genes for active (*notch1*/*pcna*/*mki67*/*fgfr3*/*wnt5b*) or quiescent ependymal cells (*edn3*/*lrrtm4*)^58^. Glutamatergic neurons were subclusters with *slc17a7* and *slc17a6* expression set >0, while *gfap* and *gad1* expression was set =0. With a resolution of 0.8, this generated 21 cell clusters from a total of 4,960 glutamatergic neurons. Finally, GABAergic neurons were subclustered by the expression of gad1 and gad2 set >0, while slc17a6, slc17a7 and gfap were set =0. At a resolution of 0.8, this subclustered 3,392 GABAergic neurons and generated 18 cell clusters.

Differential gene expression was run to determine the log2fold change between specific cell clusters in each treatment group. The log2fold change of candidate genes identified using GOrilla software were plotted on a heatmap using GraphPad Prism software.

### Gene Ontology Analysis

Gene Ontology (GO) terms were determined using GOrilla software. We included the top differentially expressed genes, calculated using Seurat, with the highest log2fold change in the GOrilla analyses. GOrilla generated a list of enriched biological processes, and we selected candidate genes based on their log2fold expression.

### Real Time Qualitative PCR

The hypothalamus was microdissected from larval (2-3cm) axolotl animals at 1 day post tail or limb amputation and from uninjured control animals, then immediately flash frozen on liquid nitrogen. Moreover, hypothalamus tissue was also collected at 1-day post tail amputation from SR142948A-injected and vehicle control animals. In all experiments, five hypothalamus regions were pooled for each biological replicate, and three biological replicates were used in all experiments. Finally, five tail tissues were pooled from control or SR142948A-injected animals at 1-day post tail amputation, and a total of three biological replicates were utilized in all experiments. Total RNA was extracted using Trizol Reagent according to manufacturer’s instructions, and cDNA was synthesized from 1μg of total RNA using the iScript cDNA Synthesis Kit (BioRad). RT-qPCR was performed using a BioRad CFX Opus 96 detection system.

Custom primers were designed against axolotl-specific sequences, including:

*18s* For: CGGCTTAATTTGACTCAACACG,*18s* Rev: TTAGCATGCCAGAGTCTCGTTC; *gal* For: CCCGACCTCCTCAGAAGAAA, *gal* Rev: AGGCAGGTTTCTCAGTGGAA; *ghrh* For: TTCACACTGTTCCCCGCTAT, *ghrh* Rev: TCTCGACTCTGCACTGGATC; *cck* For: GAGCTCTTCTGGACAGCAGA, *cck* Rev: TCTGGCCACTAAAGCTCCAA; *prlh* For: TCTTCAGGAACGTGGAGTCC, *prlh* Rev: AGCAACGTTACTGTGCACTG; *il1b* For: AAAGGATATTCACACCGCGC, *il1b* Rev: CACGGGTAGAAGTTGTGTGC; *il8* For: GTTCCACCCCAAACGAGAAC, *il8* Rev: GCCAACTCTTCATTGCCCAA.

### Statistical Analyses

All statistical analyses were performed using GraphPad Prism Software v10. Analyses included t-tests to compare two treatment groups, or a one- or two-way ANOVA with a Tukey post hoc test for multiple comparisons. Differences between groups were considered significant at four different levels, including: *p<0.05, **p<0.01, ***p<0.001, ****p<0.0001.

## Data Availability

Raw sequencing data is deposited on the NCBI SRA under BioSample accessions: SAMN45909195,SAMN45909196,SAMN45909197 and SAMN45909198

## Acknowledgements

This work was supported by funding from the Owens Family Foundation and start-up funds from the MBL. S.E.W. was supported by a NSERC PDF fellowship.

## Figure Legends

**Supplementary Figure 1. Erk activation in the brain after tail amputation is maintained into adulthood. A)** Schematic depicts the approximate location of cross sections throughout the telencephalon. **B, C)** Immunostaining using a dpErk antibody on brain tissue 1-hour post tail amputation in larval (B) and adult animals (C). **D, E)** Immunostaining using a dpErk antibody on axolotl brain tissue at 3-days post tail amputation in both larval (D) and adult (E) animals. Scale bars: 500μm.

**Supplementary Figure 2. Erk activation in the brain in response to different injuries**. Immunostaining for dpErk and NeuN in the axolotl brain at 1-day post spinal cord injury (A), limb amputation (B), and a skin punch biopsy (C). Scale bars: 200μm. Scale bars for iii images: 100μm.

**Supplementary Figure 3. pAAV viral tracing of neurons within the medial pallium**. Serial vibratome sections of an entire axolotl brain demonstrate the labelling of cells exclusively within the medial pallium and their corresponding axonal projections. Scale bars: 200μm.

**Supplementary Figure 4. Subclustering of ependymoglia, glutamatergic neurons and GABAergic neurons. A)** UMAP of 18 ependymoglia subclusters, labelled as quiescent or active. **B)** Feature plots demonstrate the expression of genes associated with active or quiescent ependymoglia. **C)** Dotplot demonstrates the expression of marker genes for active or quiescent ependymoglia in each cluster. **D)** UMAP of 18 ependymoglia subclusters labelled based on their cell fate. **E)** Dotplot demonstrates the expression of marker genes for glutamatergic or GABAergic neurons, along with oligodendrocytes and choroid plexus epithelial cells in each ependymoglia cell cluster. Ependymoglia expressing markers for these cell types were labelled as progenitor cells. Green and blue boxes indicate whether these cell clusters were originally identified as active or quiescent. **F)** UMAP displays 21 glutamatergic cell clusters. **G)** Feature plots for glutamatergic marker genes. **H)** Dotplot demonstrates the expression of glutamatergic marker genes and the absence of ependymoglia or GABAergic neuron markers. **I)** UMAP displays 18 GABAergic neuron clusters. **J)** Feature plots demonstrate the expression of GABAergic marker genes. **K)** Dotplot demonstrates the expression of GABAergic marker genes across each cell clusters, and the absence or glutamatergic neuron and ependymoglia markers.

**Supplementary Figure 5. Conservation of mammalian marker genes in axolotl *etv1*^+^/*slc17a7*^+^ neuron clusters. A)** Dotplot demonstrates the expression of mammalian marker genes for thalamus-projecting neurons born in the cortex at various stages of mouse development. **B)** Dotplot shows the expression of mammalian cortex-projecting neurons born in the cortex that are found in various stages of mouse development. **C)** Dotplot shows the expression of mammalian marker genes for hippocampal-born neurons, including general marker genes and those for specific regions of the hippocampus (granule, CA1, CA2, CA3).

