## Supplementary figures and images for "Neuronal activation in the axolotl brain promotes spinal cord regeneration"

### Supplemental Figure 3

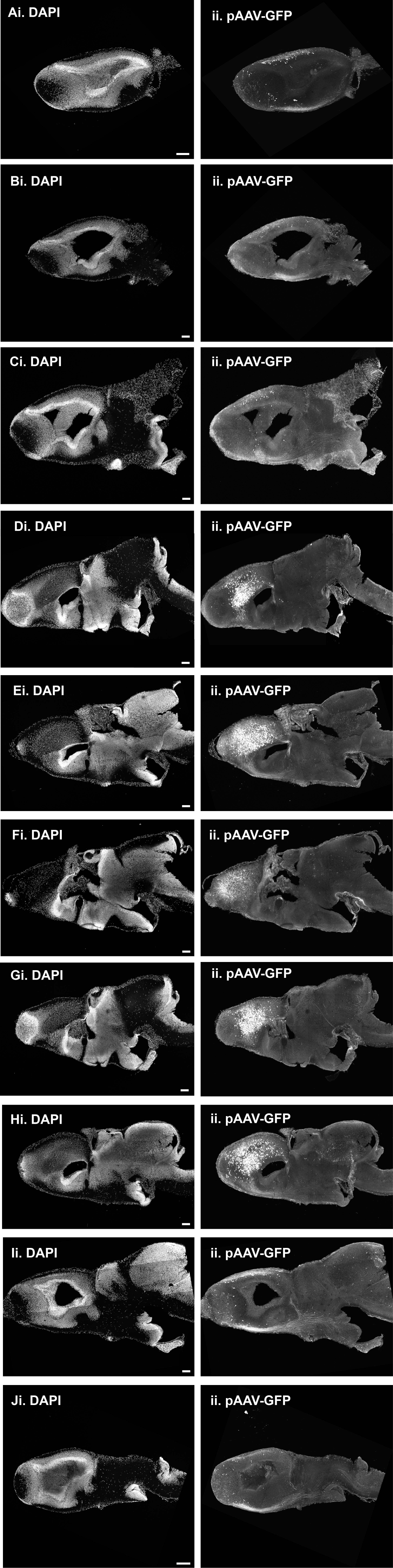

### Supplemental Figure 4

Supplementary Figure 4

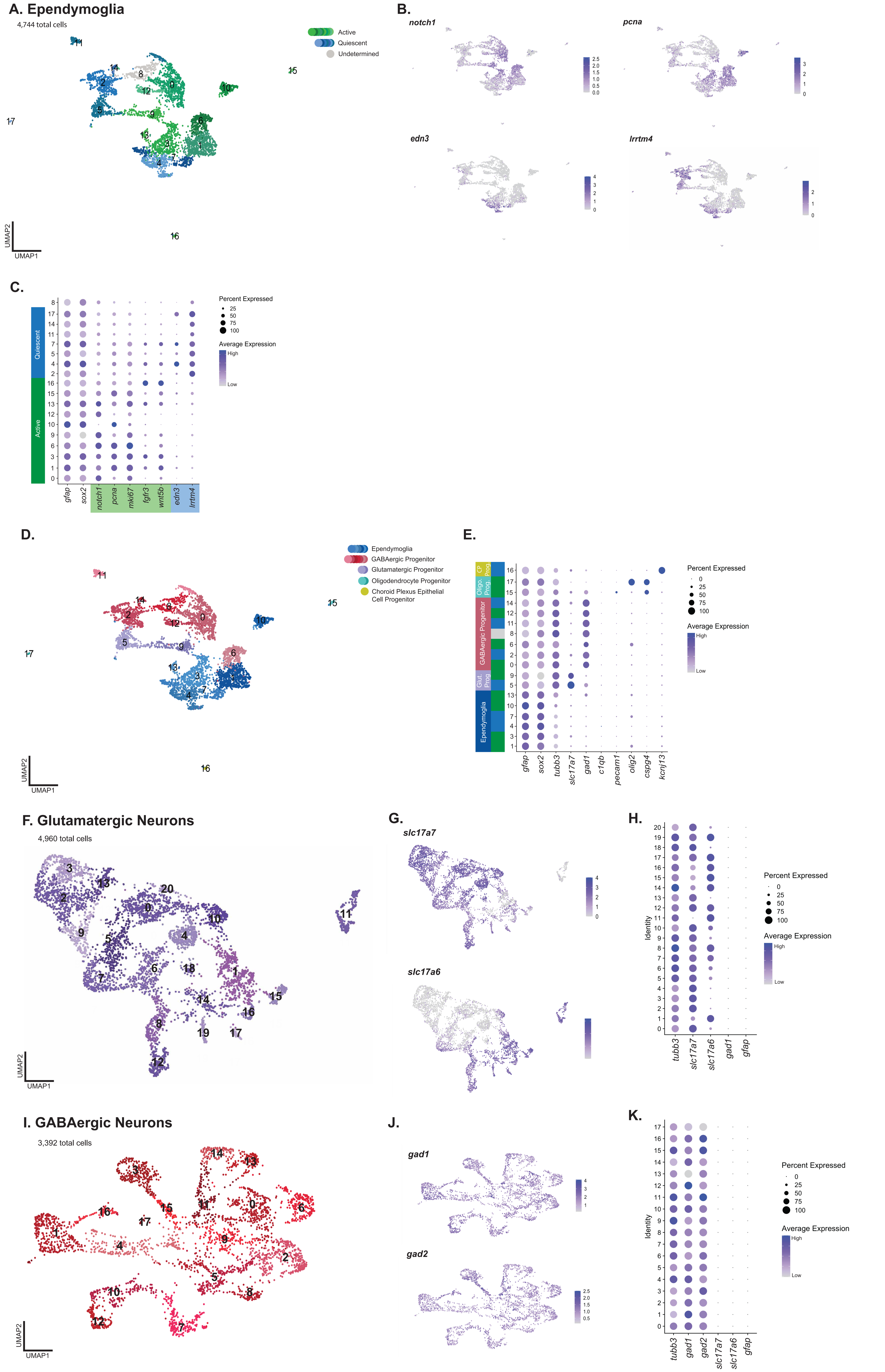

### Supplemental Figure 5

#### Supplementary Figure 5

##### A. Cortex Neurons

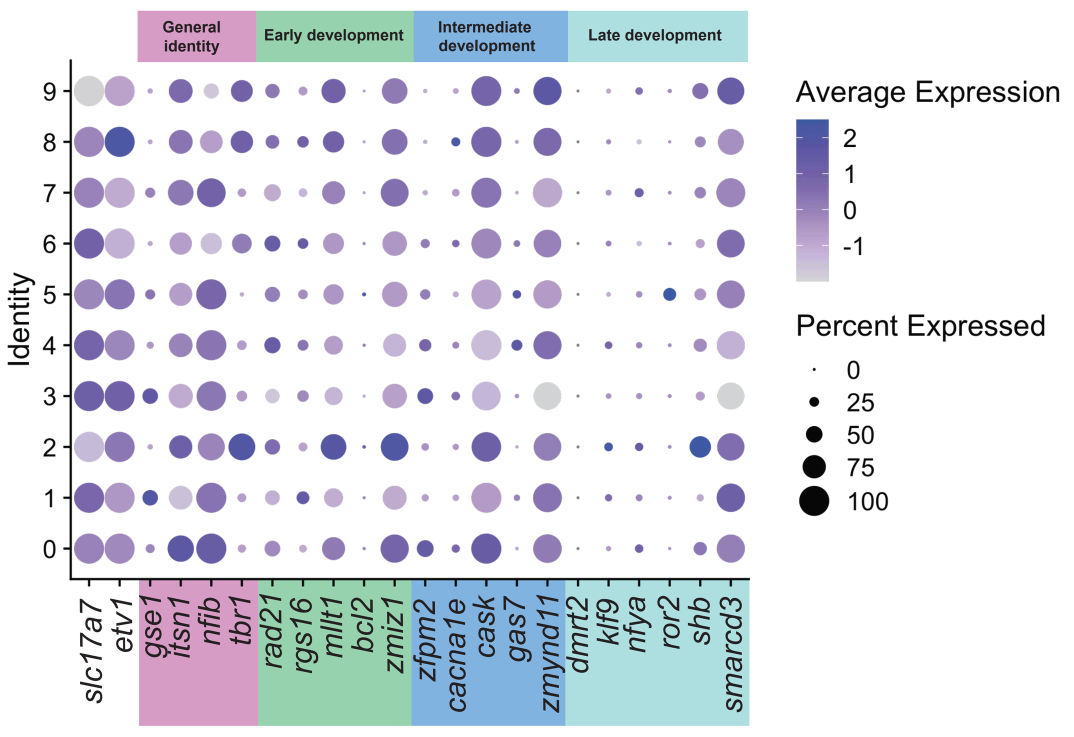


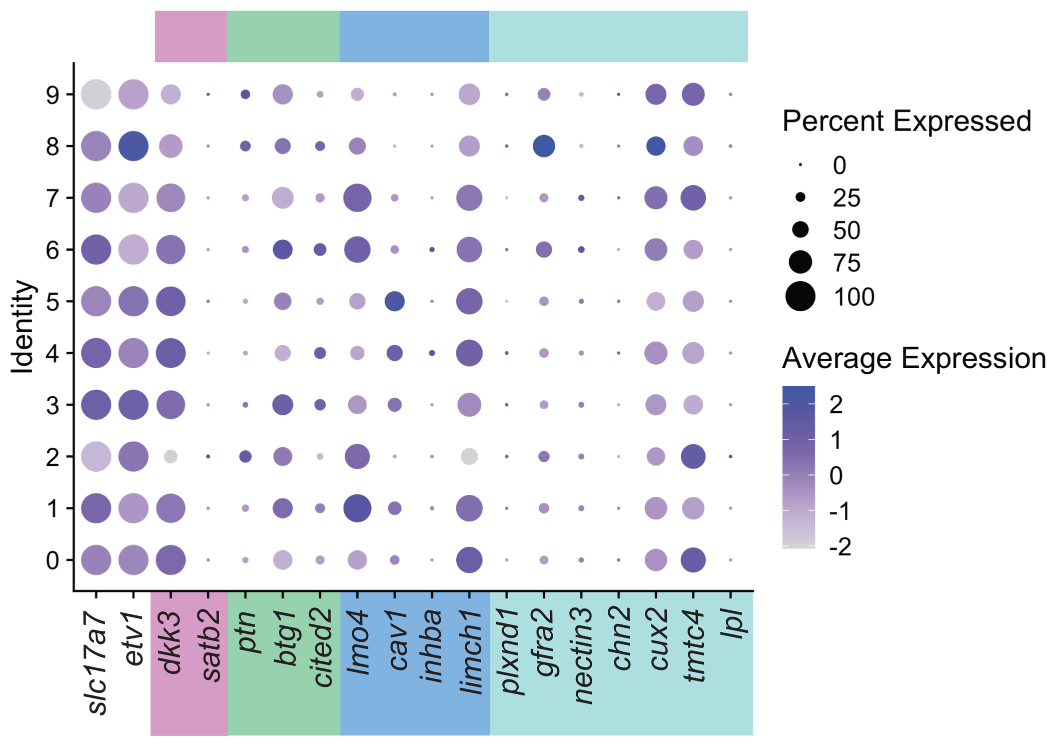


##### B. Hippocampal Neurons

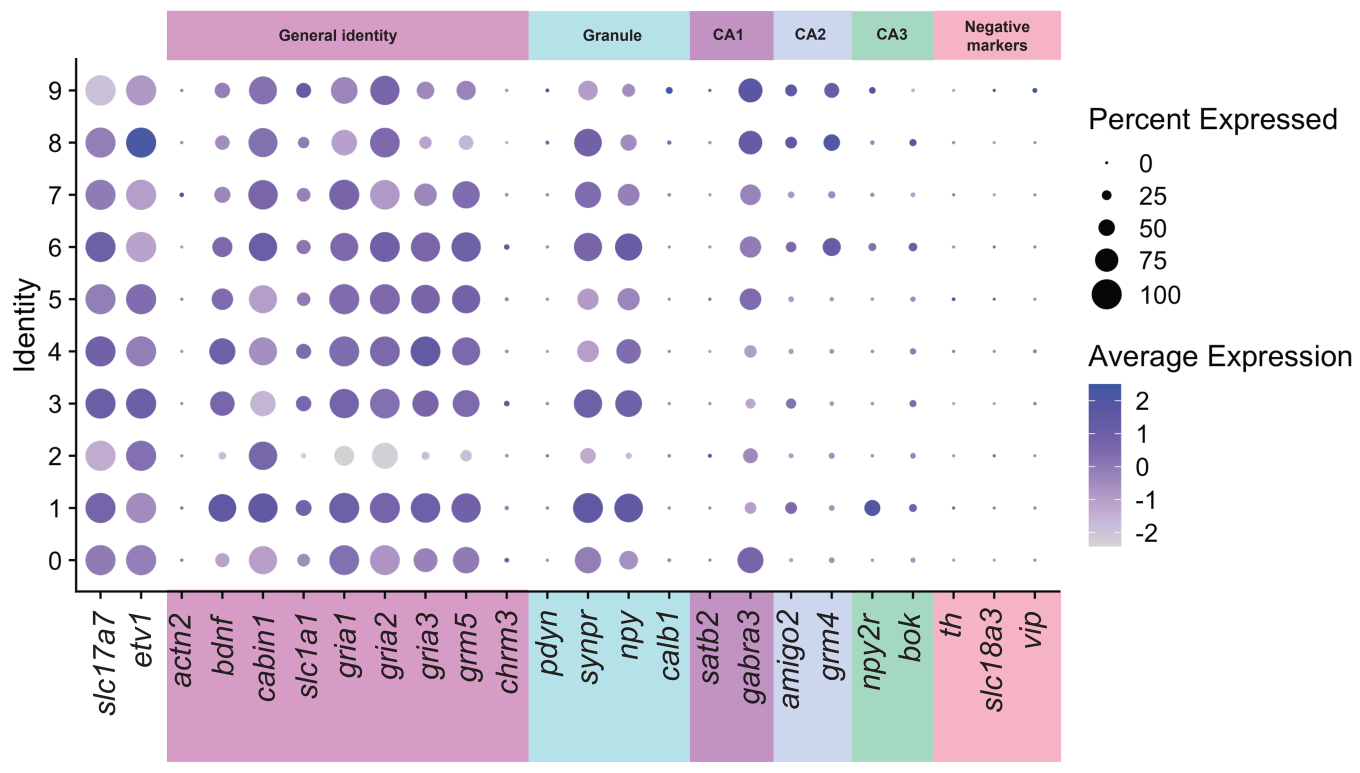

### Supplementary Figure 1

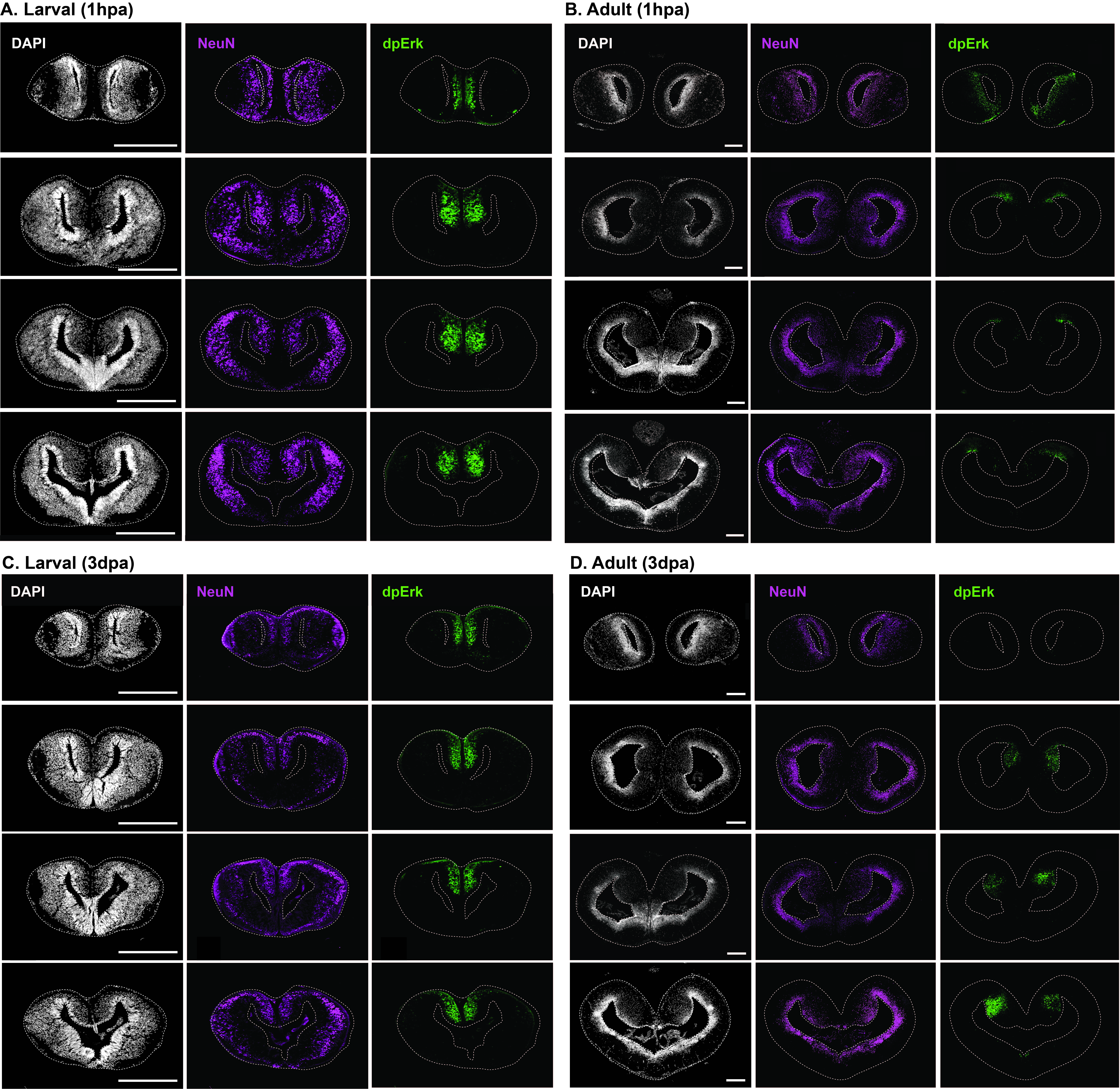

### Supplementary Figure 2

#### Supplementary Figure 2

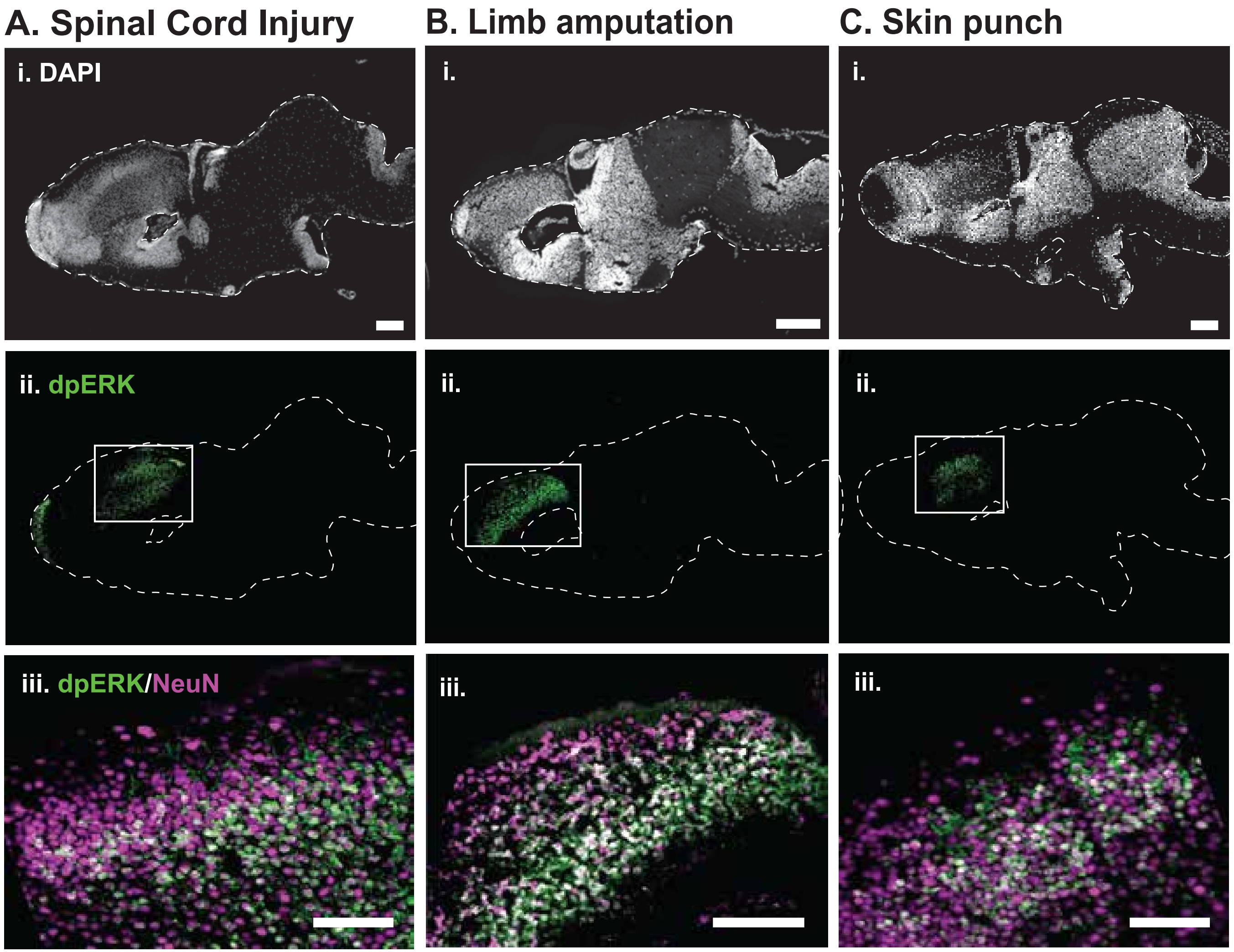
